# A Simple and Scalable Chopped-Thallus Transformation Method for *Marchantia polymorpha*

**DOI:** 10.1101/2025.01.18.633702

**Authors:** Rituraj Batth, Andisheh Poormassalehgoo, Kritika Bhardwaj, Elżbieta Kaniecka, Shino Goto-Yamada

## Abstract

The liverwort *Marchantia polymorpha* has emerged as a valuable model for studying fundamental biological processes and the evolutionary history of land plants. Agrobacterium-mediated transformation is widely used for genetic modification of *M. polymorpha* using spores, thalli, and gemmae. While spores offer high transformation efficiency, they result in diverse genetic backgrounds due to sexual reproduction. Conversely, thallus- and gemma-based methods maintain genetic consistency but are impractical for large-scale applications. To address these limitations, we developed a novel chopped-thallus transformation method. This technique involves generating numerous plant fragments by chopping thalli, eliminating the need for complex preprocessing. The method demonstrated superior transformation efficiency compared to traditional approaches and achieved sufficient numbers of transformants using simplified Gamborg’s B5 medium, previously considered suboptimal. This scalable and straightforward method enables the generation of large numbers of genetically consistent transformants, facilitating high-throughput experiments, including mutant screening and other large-scale applications.

## 1. Introduction

The liverwort *Marchantia polymorpha* (hereafter Marchantia) belongs to one of the earliest diverging lineages of land plants, along with hornworts and mosses [1,2]. Its genome has been fully sequenced, and the data are publicly available. Marchantia’s simple morphology and compact structure make it particularly suitable for laboratory studies [1,3,4]. In addition to sexual reproduction, Marchantia can propagate asexually through detached thalli or gemmae, which is beneficial for analyzing mutants with reproductive abnormalities. Moreover, its haploid-dominant life cycle and low level of gene redundancy have facilitated its use in genetic studies [1,3,4]. For these reasons, Marchantia has emerged as a key model organism in plant research.

Genetic modification is an essential technique for studying molecular biology, with significant implications for agriculture, biotechnology, and fundamental biological research. Several methods have been proposed for transforming Marchantia, including Agrobacterium-mediated stable and transient transformations [5-8] and particle bombardment [9]. Agrobacterium-mediated transformation is widely used approach due to its efficiency in stably integrating foreign DNA into plant genomes. In Marchantia, Agrobacterium-mediated transformations have been established using spore-derived tissues, thalli, and gemmae to obtain stable transformants. Thallus-based transformations involve excising apical meristems, regenerating fragments on plates, and co-cultivating them with Agrobacterium in immersion culture [6]. The spore-based method is the most efficient to date, where spores germinate into immature thalli, which are then co-cultivated with Agrobacterium [5]. The AgarTrap method, which combines plant culture, infection and selection on a single plate, has also been developed [7,10].

Although Agrobacterium-mediated transformation is widely used in Marchantia species, several challenges remain unresolved. First, transformation efficiency can vary significantly, and there is a need for methods that consistently improve this efficiency. Second, traditional transformation methods often rely on complex media compositions [5,6], therefore, evaluating the efficiency of simpler, more accessible media would be highly beneficial. Moreover, obtaining a large number of transformants, such as those used for mutant screening, is crucial for large-scale experiments. The most efficient spore-based method presents challenges: spores generated from the mating of male and female strains can lead to genetically heterogeneous populations. While the male Takaragaike-1 (Tak-1) and female Tak-2 strains share highly similar traits, they differ in reproductive chromosomes, numerous SNPs, and subtle morphological variations [1,5,11]. Recent studies have also reported differences in stress responses between male and female plants [12]. Although the Tak-1 inbred female line BC3-38 has been established from an F1 hybrid of Tak-1 and Tak-2 [5,13], it suffers from low fertilization success. Although spore-based transformation remains the most efficient method, the use of asexually propagated thalli is preferred due to these genetic considerations. Additionally, mutants with reproductive defects, such as autophagy mutants [14], are unable to produce spores from their genetic background. These limitations highlight the need for a simplified, efficient transformation method utilizing vegetatively propagated tissues, such as regenerating thalli.

Marchantia, like other bryophytes, has high regenerative ability, allowing for the easy establishment of new plants [15,16]. Removing the meristem can induce the regeneration of thallus edges, leading to the formation of new tissues within five days [6]. Protoplast cells in Marchantia can also regenerate their cell walls and reproduce a whole plant [17,18], suggesting the potential for regenerating entire plants from single cells. In traditional thallus transformation methods, one Marchantia plant produces four thallus fragments for subsequent transformation [6]; however, fragments can potentially be smaller, with studies reporting of transformations using fragments as small as 4–5 mm square [19]. In the amphibious liverwort *Riccia fluitans* and the hornwort *Anthoceros agrestis*, Agrobacterium-mediated transformation has been performed with fragmented thalli obtained by chopping [20-22].

As previously mentioned, several challenges persist, including inconsistent transformation efficiencies, the reliance on complex media compositions, and difficulties in producing large numbers of genetically consistent transformants. To address these issues, we developed an optimized Agrobacterium-mediated transformation protocol using finely chopped thallus fragments, which takes advantage of the remarkable regenerative capacity of Marchantia. We developed a scalable method that generates numerous transformants from minimal starting material. We evaluated factors influencing transformation efficiency, such as regeneration duration and medium composition. This method offers a solution for the genetic modification of Marchantia, enabling large-scale applications, including mutant screening and other high-throughput studies.

## 2. Materials and Methods

### 2.1 Plant material and preparation

The male accession of *Marchantia polymorpha* Takaragaike-1 (Tak-1) [5,23] was kindly provided by Dr. Takayuki Kohchi (Kyoto University, Japan) and Dr. Shoji Mano (National Institute for Basic Biology, Japan). Plants cultured on a solid ½ Gamborg’s B5 medium (half-strength Gamborg’s B5 [Metck, G5768] with 2.5 mM MES-KOH [pH 5.7]) supplemented with 1% (w/v) sucrose and 1% (w/v) phytoagar (Duchefa Biochemie, P1003). Unless otherwise specified, the plants were grown under continuous white, fluorescent light at an intensity of 50 µmol m^−2^ s^−1^ at 22°C.

### 2.2 Analysis of thallus size and regeneration

Marchantia gemmae were grown on ½ Gamborg’s B5 medium plates supplemented with 1% sucrose for 12 days. The thalli were then chopped into fragments smaller than 2 mm using a sterilized razor blade and washed four times with growth medium. The fragments were spread onto ½ Gamborg’s B5 medium with 1% sucrose plates and incubated at 22°C under continuous white, fluorescent light for 7 days. Images of 16 plate sections were captured using a stereomicroscope on days 0 and 7 to assess survival and growth. The survival rate was determined based on the presence of viable green tissue and the visible regeneration of new tissue. The size of the thallus fragments was measured using Fiji/ImageJ software (https://imagej.net).

### 2.3 Plasmid construction

The reporter gene *β-glucuronidase* (*uidA*/*GUS*) expressing construct (EF1pro:GUS) was generated as follows. The *GUS* gene was amplified with the primers GUS_fuF: 5’-AAGGAACCAATTCAGTCGACATGTTACGTCCTGTAGAAACCCC-3’ and GUS_fuR: 5’-AAGAAAGCTGGGTCTAGATATCATTGTTTGCCTCCCTGC-3’. The amplified fragment was cloned into the *Sal*I/*EcoR*V-digested pENTR1A vector (Thermo Fisher) via Gibson assembly. The *GUS* gene was subsequently transferred into the pMpGWB303 plasmid [24] using Gateway LR cloning (Thermo Fisher). This construct enables the expression of GUS under the control of the Marchantia elongation factor 1-α (EF1α) promoter.

### 2.4 Bacterial strain and culturing

Agrobacterium, *Agrobacterium tumefaciens* strain EHA101, carrying the EF1pro:GUS plasmid was used in all experiments. The EH101 strain has high transformation efficiency in Marchantia transformation [7]. For transformation, Agrobacterium cultures were initiated from a single colony and grown in 5 mL of LB medium at 30°C with overnight agitation. One millimeter of the bacterial culture was centrifuged to pellet the cells, which were then resuspened in 2.5 ml of ½ Gamborg’s B5 medium supplemented with 1% (w/v) sucrose and 100 µM 3,5-dimethoxy-4-hydroxyacetophenone (acetosyringone). The suspension was incubated at 30°C for 3–6 hours to induce virulence before co-cultivation with Marchantia thallus fragments. All procedures were performed aceptically.

### 2.5 Agrobacterium-mediated transformation of Marchantia

For the chopped-thallus method, a total of ten 12-to 14-day-old Marchantia plants were collected and placed in a plastic Petri dish with 3–5 mL of liquid medium. Using a sterilized razor blade, the thalli were finely chopped into fragments smaller than 2 mm within a laminar flow cabinet. The fragments were washed three times with liquid medium using a cell strainer. The chopped thallus pieces were cultured in 50 mL of ½ Gamborg’s B5 medium with 1% sucrose or a supplemented medium containing 1% sucrose, 0.03% L-glutamine, and 0.1% of Casamino acid in a 250-mL flask capped with aluminum foil. Cultures were incubated under continuous white fluorescent light (50 µmol m^−2^ s^−1^) at 22°C for 3–7 days with agitation at 180 rpm on a shaker to promote regeneration. Before co-cultivation, an additional 2.5 mL of 20% sucrose (equivalent to 1% sucrose in the total volume) was added to the flasks containing medium with 1% sucrose. While this brought the theoretical total sucrose concentration to 2%, the actual concentration may have been lower due to the consumption of sucrose by the thallus fragments during regeneration. One milliliter of Agrobacterium culture (prepared as described in Section 2.4) and 100 µM acetosyringone were then added. Co-cultivation was performed with agitation at 180 rpm at 22°C in the dark for 3 days. Following co-cultivation, the fragments were washed with sterilized water four times as follows: The culture from a flask was transferred to a Falcon tube and allowed to settle briefly, enabling the fragments to sediment. The supernatant was carefully removed as much as possible, and sterilized water was added to the tube. The contents were shaken or vortexed, and the supernatant was removed as described above. Sterilized water was added again, and the contents were shaken or vortexed before being poured through a cell strainer. The fragments were subsequently rinsed twice with sterilized water. The washed fragments were transferred to selection plates containing ½ Gamborg’s B5 medium supplemented with 0.5 µM chlorsulfuron and 100 mg/L cefotaxime and incubated under continuous white light at 22°C.

For the traditional thallus method [6], ten 12-to 14-day-old Marchantia plants were harvested, and their apical meristems were removed. Each thallus was divided into two pieces, resulting in four pieces per plant. These fragments were cultured on a solid ½ Gamborg’s B5 medium with 1% sucrose and 1% agar under continuous light at 22°C for 3 days to promote regeneration. Forty regenerating plantlets were co-cultivated with 1 mL of Agrobacterium culture (prepared as described in Section 2.4) in 50 mL of ½ Gamborg’s B5 medium containing 2% sucrose. Additionally, co-cultivation was performed with medium supplemented with 0.03% L-glutamine and 0.1% casamino acid in a 250-mL flask. Both cultures were supplemented with 100 µM acetosyringone. The subsequent washing and selection steps were identical to those used for the chopped-thallus method.

### 2.6 GUS enzyme activity check via histochemical assay

Histochemical assays for GUS enzyme activity were conducted with slight modifications based on previously described methods [25]. Plant tissues were vacuum-infiltrated with GUS assay solution for 30 min and incubated for up to 2 hours at 37°C. The GUS assay solution contained 100 mM sodium phosphate buffer (pH 7.0), 0.5 mM potassium ferrocyanide, 0.5 mM potassium ferricyanide, 10 mM EDTA, 0.01% (v/v) Triton X-100, and 1 mM 5-bromo-4-chloro-3-indolyl-β-glucuronide (X-Glc). Ethanol was used to clear pigments, ensuring better visualization of GUS staining.

### 2.7 Genomic DNA isolation and PCR analysis

Genomic DNA was extracted from transformed Marchantia to confirm the integration of T-DNA. Approximately 50 mg of thallus tissue was homogenized in a microcentrifuge tube with 50 µl of DNA extraction buffer (200 mM Tris-HCL [pH 7.5], 250 mM NaCl, 25 mM EDTA, and 0.5% [w/v] SDS). The homogenized sample was mixed with 100 µl of absolute ethanol, vortexed, and centrifuged at maximum speed for 5 min at 22°C. The resulting pellet was vacuum-dried and dissolved in Tris-EDTA buffer (pH 8.0). For PCR analysis, the extracted genomic DNA was amplified using the primers: GUS_FP (5’-ATACCGAAAGGTTGGGCAGG-3’) and GUS_RP (5’-TCTTGCCGTTTTCGTCGGTA-3’) to detect the integration of the GUS transgene. Y chromosome-specific region rhm12 [26] was amplified as a positive control using primers: rhm12_for (5’-GAGAGTATTTGCGATGCGTCAC-3’) and rhm12_rev (5’-CAAGGGCTCGAATCCATTTCT-3’).

### 2.8 Statistical analysis

Statistical analysis was conducted using R software (version 3.2.1; R Core Team, 2025). Data were analyzed using one-way ANOVA followed by Tukey’s HSD test at a 0.05 significance level.

## 3. Results and Discussion

### 3.1 Thallus Finely Chopped into Micro-Fragments Generates a Large Number of Regenerated Plants

As previously reported, excised pieces of thallus produce visible tissue five days after plating on growth medium, suggesting that regenerating thalli undergo extensive cell division without the application of growth regulators [6,27]. Fragments traditionally used in thallus transformation are approximately 5–8 mm in size. This study aimed to determine whether smaller thallus fragments could regenerate effectively.

To investigate this, thalli, which have developed from gemmae and 12-day-old plants with dichotomous branches and meristems at each growth tip, were chopped into fragments smaller than 2 mm using a sterilized razor blade (Figure 1A). The fragments were placed on a solid growth medium and incubated under standard growth conditions. The fragmented thalli exhibited substantial regenerative capacity, with 88% of the fragments surviving. Even fragments as small as 0.2 mm regenerated and formed new tissues (Figure 1B). Survival rates varied with fragment size: fragments measuring 0.19–0.27 mm (0.035–0.071 mm^2^) achieved over 70% survival, while fragments larger than 0.49 mm consistently demonstrated 100% survival and regeneration (Figure 1C).

**Figure 1.**
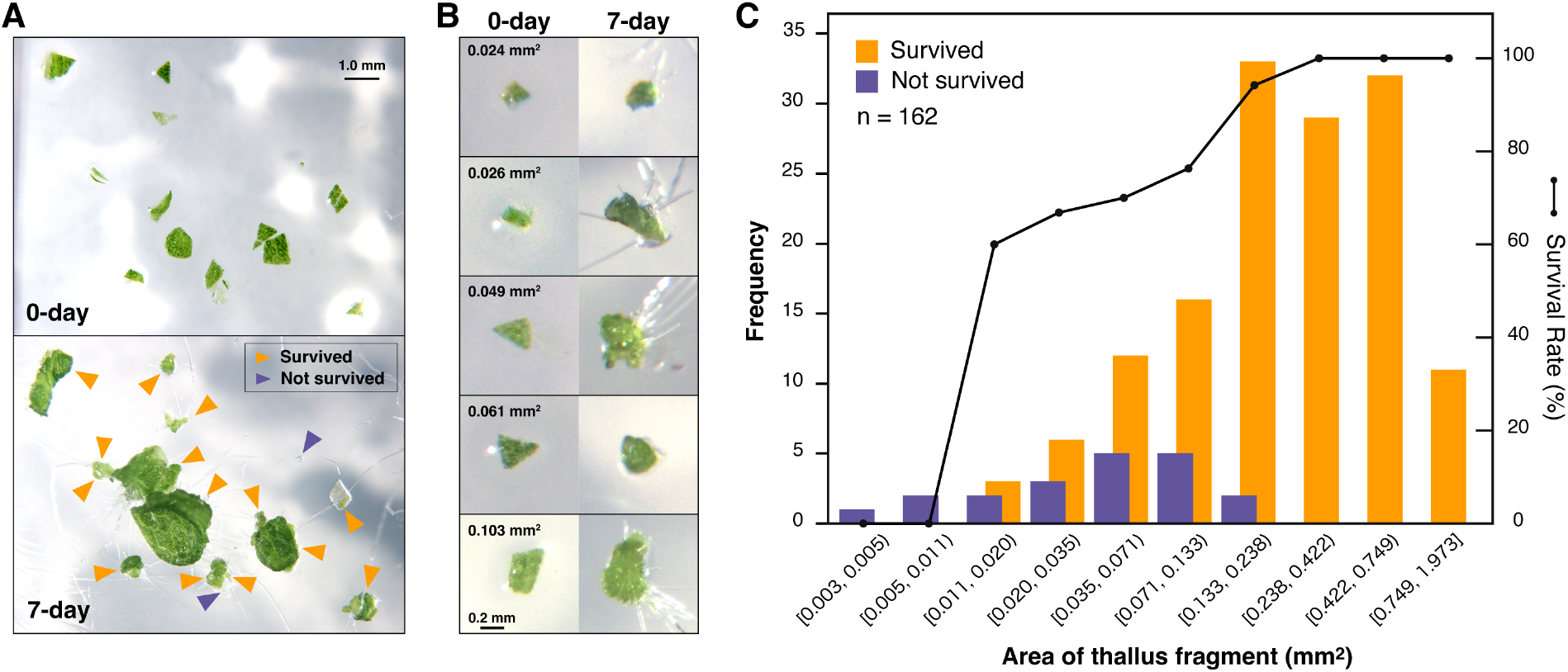
Survival rate of chopped thalli. (A) Whole Marchantia plants, aged 12 days, were finely chopped into fragments smaller than 2 mm using a razor blade. These fragments were incubated on growth medium plates at 22°C under continuous light for 7 days, and their growth was observed under a microscope. Survival was determined by green coloration and visible cell proliferation. (B) Magnified images of surviving thallus fragments. The numbers in the top-left corner represent the area of each fragment. (C) A histogram showing the number of surviving and dead thallus fragments based on their size. The survival rate for each size category is represented as a line graph derived from this data. The data were binned using logarithmic scaling. A total of 162 thallus fragments were analyzed.

This study demonstrates that the chopped thallus fragments can regenerate, highlighting the robustness of Marchantia in tissue regeneration. The failure of smaller fragments to regenerate may be attributed to the absence of cells capable of regeneration, potentially due to physical damage caused by chopping or a decline in cell activity for some unknown reason. Larger fragments are more likely to contain intact and viable cells, which may explain their higher regeneration rates. Nonetheless, it is worth emphasizing that most fragmented thallus pieces, except for the extremely small ones, regenerated successfully. Chopping generates a large number of viable and regenerating tissues from minimal initial material, providing a scalable and practical solution for high-throughput applications.

### 3.2 Establishing the Chopped-Thallus Method for Thallus Transformation

In spore-based Marchantia transformations, immature thalli derived from spores are typically used [5,10]. The chopped-thallus method proposed in this study generates fragments smaller than those traditionally used in thallus-based transformations, yet they remain suitable for genetic modification. To evaluate the transformation efficiency of this method, we followed the protocol described by Kubota et al., 2013 [6]. Hereafter, we refer to the chopped-thallus approach as the “chopped-thallus method” and the established approach as the “traditional method.”

Thalli grown on a solid medium for 12–14 days from gemmae served as the starting material for transformation (Figure 2A). For the chop-thallus method, the thalli were finely chopped into fragments (<2 mm) using a razor blade (Figure 2B). Regeneration of these fragments was performed in liquid ½ Gamborg’s B5 medium supplemented with 1% sucrose (Figure 2C), allowing co-cultivation with Agrobacterium to proceed in the same container, eliminating the need to transfer the fragments. The fragments were co-cultivated with Agrobacterium carrying reporter genes (GUS and chlorsulfuron-resistant gene) (Figure 2D). Sucrose is essential for successful transformation [6]. In this study, 1% sucrose was added to the co-cultivation medium, resulting in a total of 2% sucrose during co-cultivation, accounting for the sucrose already present. Acetosyringone was also added to enhance Agrobacterium virulence. In the traditional method, plants were processed by removing apical meristems and dividing the thalli into four pieces, followed by regeneration on solid growth medium (Figure 2F,G). These regenerated fragments were co-cultivated with Agrobacterium in ½ Gamborg’s B5 medium containing 2% sucrose and acetosyringone under dark conditions (Figure 2H).

**Figure 2.**
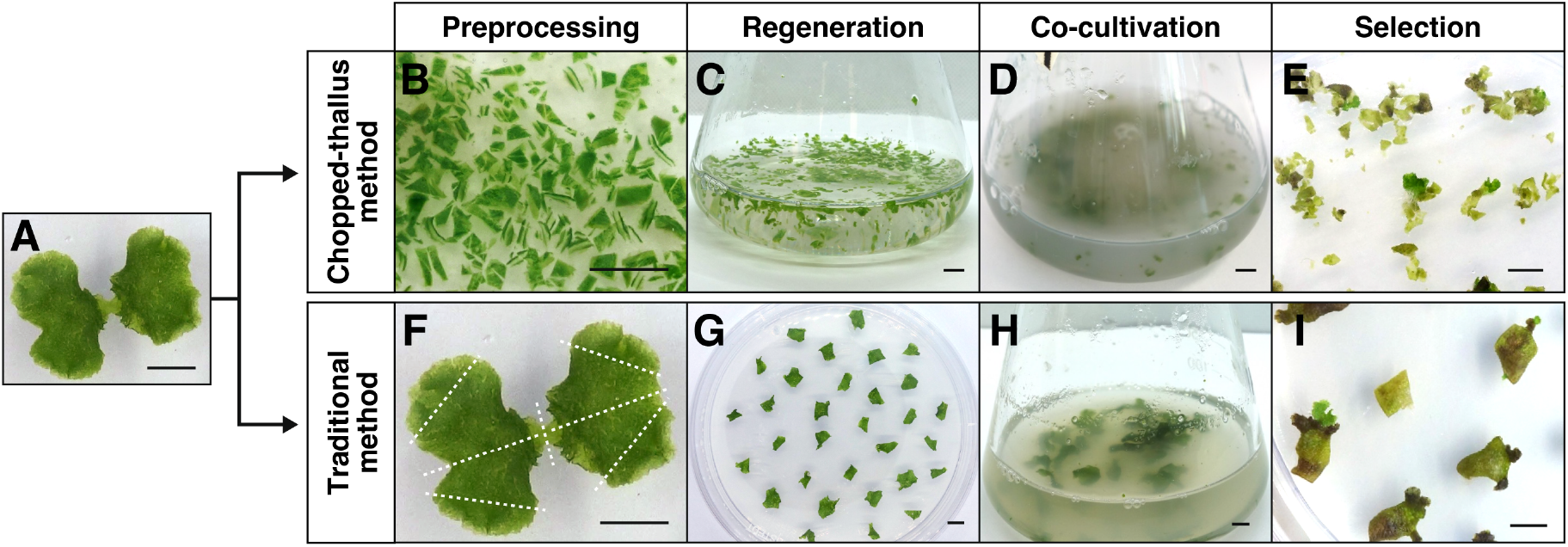
Workflow of thallus transformation. Marchantia plants aged 12–14 days (A) were used as materials for transformation using either the chopped-thallus method (B–E) or the traditional method (F–I). Bars = 5 mm.

Selection with chlorsulfuron in both methods resulted in the formation of resistant plantlets after 2–3 weeks. Fresh green-resistant plantlets emerged from the edges of the basal tissue, while the original tissue turned yellow and eventually became necrotic (Figure 2E,I). Surviving plants were transferred to a new selection medium and grown for an additional week. Transformation efficiency was assessed by histochemical GUS staining. The integration of transgenes was confirmed through PCR analysis of genomic DNA from randomly selected plants (Figure 3).

**Figure 3.**
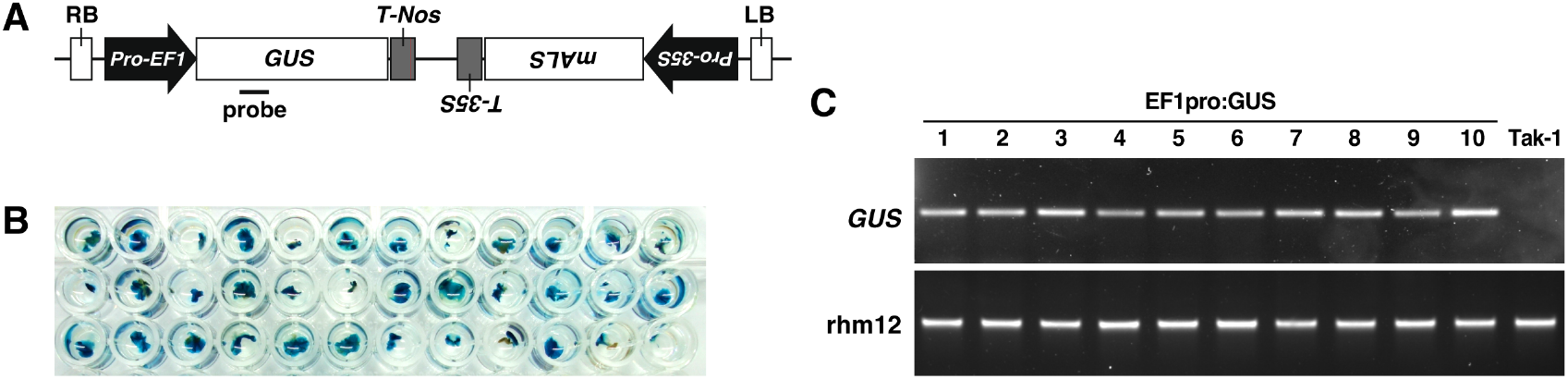
Confirmation of transformation. (A) Schematic representation of the EF1pro:GUS construct. The probe region indicates the primer binding site for the GUS gene. GUS: β-glucuronidase; mALS: mutated acetolactate synthase (chlorsulfuron resistance gene); Pro-EF1: promoter of Marchantia elongation factor 1-α; Pro-35S: cauliflower mosaic virus 35S promoter; T-Nos: nopaline synthase terminator; T-35S: cauliflower mosaic virus 35S terminator; RB: Right border; LB: Left border. (B) Transgene expression was confirmed through GUS staining of chlorsulfuron-resistant plants. (C) PCR analysis was performed on genomic DNA extracted from randomly selected chlorsulfuron-resistant plants.

### 3.3 The Chopped-Thallus Method Simplifies Transformation and Enhances Efficiency

Kubota et al. (2013) optimized the Agrobacterium-mediated thallus transformation protocol and established a consistent and reproducible method. Their key findings included: (i) a 3-day regeneration period was sufficient for effective transformation, (ii) the presence of sucrose during co-cultivation significantly enhanced transformation efficiency, and (iii) the highest transformation efficiency was achieved using the M51C medium rather than ½ Gamborg’s B5 medium. To develop the chopped-thallus method, we adopted two critical factors: (i) a 3-day regeneration period and (ii) the addition of sucrose to the co-cultivation medium. Instead of the labor-intensive M51C medium [17], we used the simpler and more readily available ½ Gamborg’s B5 medium, leveraging the high material output generated by the chopped-thallus method.

We compared the transformation efficiencies of the chopped-thallus method and the traditional methods using a ½ Gamborg’s B5 medium with a 3-day regeneration period. Although the difference was not statistically significant, a clear trend indicated that the chopped-thallus method nearly doubled the transformation efficiency compared to the traditional method (Figures 4 and 5). The lower efficiency observed in the traditional method with ½ Gamborg’s B5 medium aligns with previous findings [6]. These results indicate that chopping increases the amount of transformation-compatible material, thus mitigating the limitations associated with using ½ Gamborg’s B5 medium.

**Figure 4.**
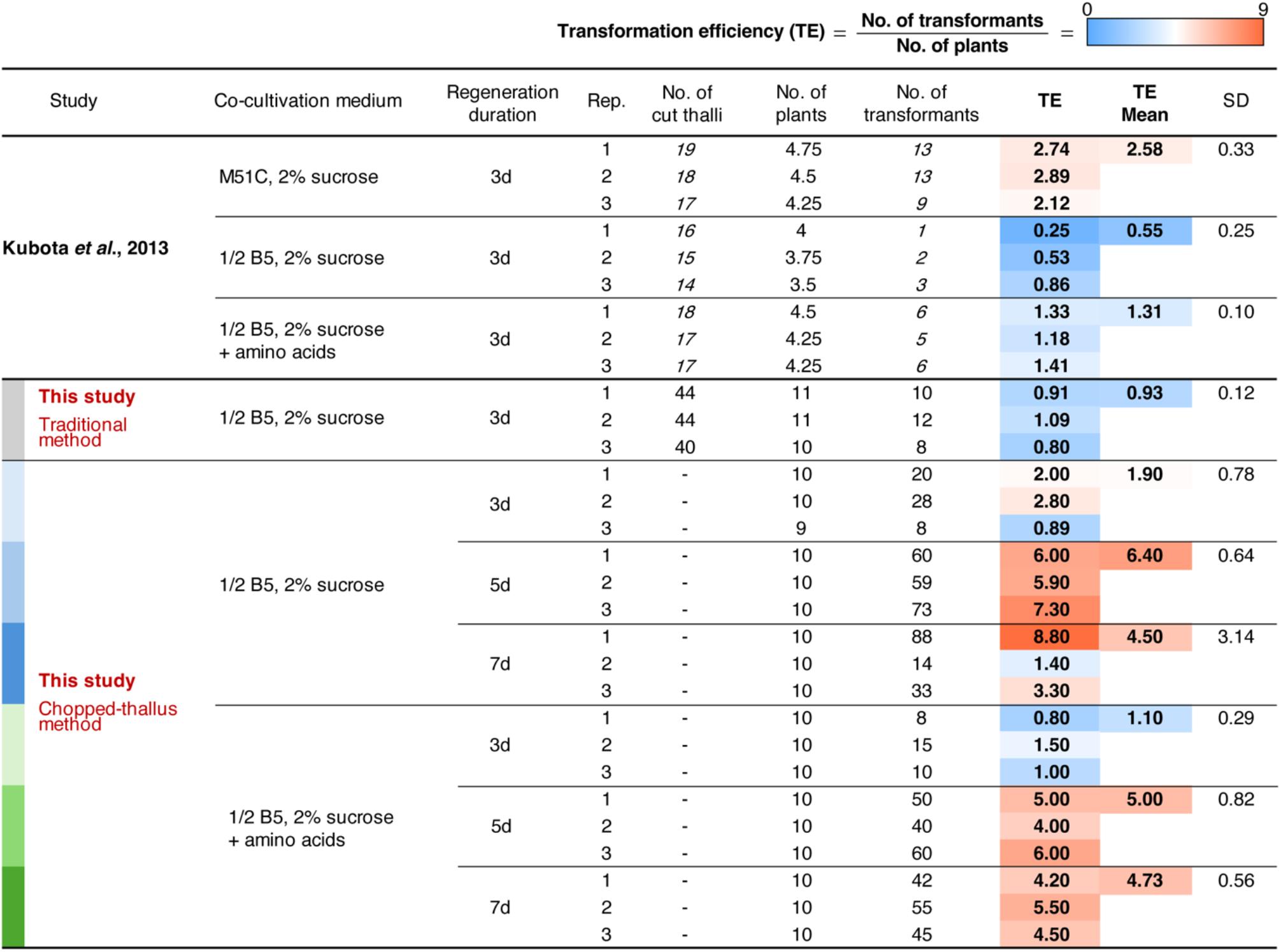
Comparison of transformation efficiency under different transformation conditions. The table summarizes the transformation efficiency for various co-cultivation media, regeneration durations, and methods (traditional and chopped-thallus). The numbers shown in italics were taken from experiments reported by Kubota et al., 2013 [6] and were used to calculate the number of plants and transformation efficiency. Experiments conducted in this study were repeated three times (Rep.). Transformation efficiency (TE) was calculated as the number of transformants per plant. The mean TE and standard deviation (SD) are shown for each condition. A heat map is applied to the TE column. The color codes on the left correspond to the charts shown in Figures 5 and 6.

**Figure 5.**
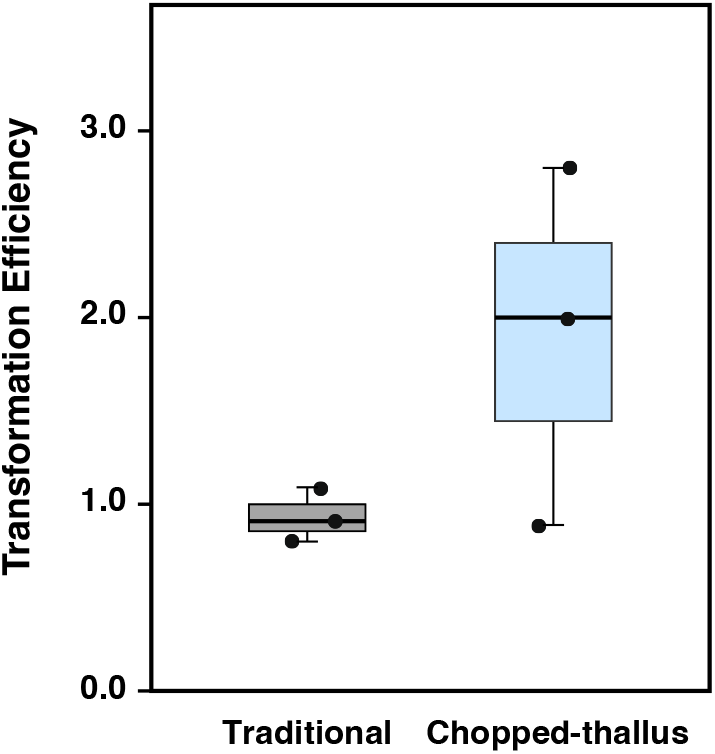
Comparison of transformation efficiency between the traditional and chopped-thallus methods. Plant fragments prepared by each method were regenerated for 3 days and then subjected to transformation. Co-cultivation was performed within ½ Gamborg’s B5 medium containing 2% sucrose. Transformation efficiency indicates the number of transformants per plant. Data represent the results of three independent experiments.

### 3.4 Optimizing Regeneration Period and Medium Composition

We further evaluated the optimal regeneration period for chopped thalli over 3–7 days, as finely chopped thalli might sustain more damage than those used in the traditional methods. Simultaneously, we investigated the effect of supplementing the co-cultivation medium with amino acids (L-glutamine and Casamino acid), which have been reported to enhance transformation efficiency [6].

Extending the regeneration period from 3 to 5 days significantly improved transformation efficiency. This trend was observed in both ½ Gamborg B5 medium and the medium supplemented with amino acids (Figures 4 and 6). In the amino acids-supplemented medium, the transformation efficiency for the 5-day regeneration period (5.00 ± 0.82) was significantly higher than for 3 days (1.10 ± 0.29). In ½ Gamborg’s B5 medium without supplementation, the transformation efficiency for 5 days (6.40 ± 0.64) showed no significant difference compared to 3 days (1.90 ± 0.78), although a potential improvement was noted.

**Figure 6.**
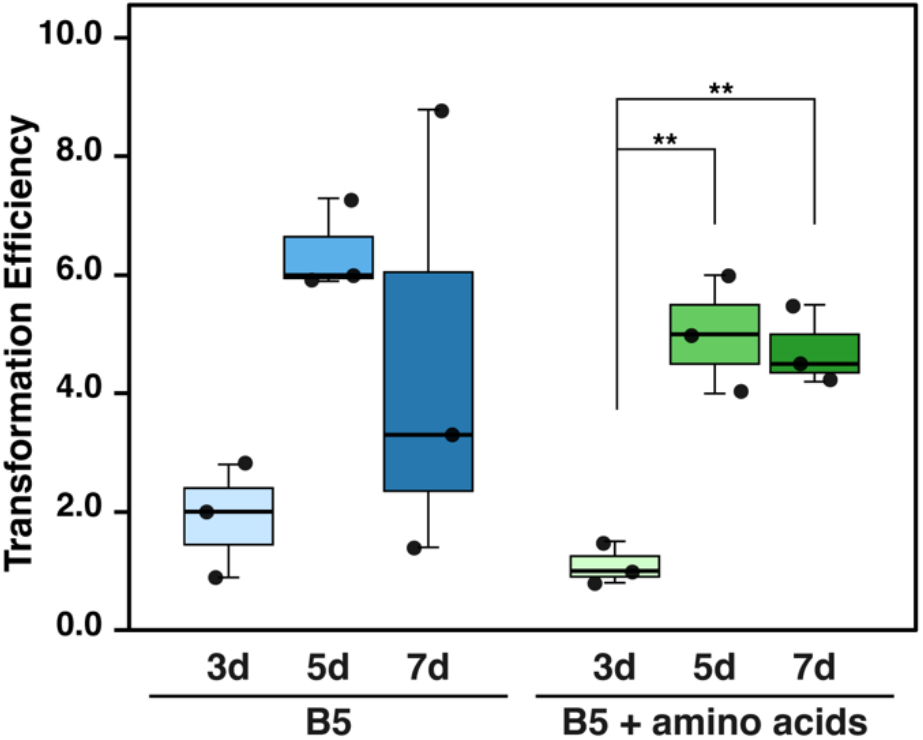
Comparison of transformation efficiency between different regeneration periods and co-cultivation media in chopped-thallus method. Regeneration periods of 3, 5, and 7 days were assessed. Two types of co-cultivation media were examined: ½ Gamborg’s B5 medium containing 2% sucrose (B5) and B5 supplemented with 0.03% L-glutamine and 0.1% Casamino acid (B5 + amino acids). Transformation efficiency indicates the number of transformants per plant. Data represent the results of three independent experiments. Statistical analysis was performed separately for each medium group using one-way ANOVA followed by Tukey’s HSD test. **p < 0.01.

Notably, the 5-day regeneration period achieved approximately 2.5 times higher transformation efficiency than the traditional method using M51C medium (2.48 ± 0.33) (Figure 4). However, no additional benefits were observed with a 7-day regeneration period compared to 5 days. Additionally, excessive growth of the thallus fragments during the 7-day period made the procedure less practical. Therefore, a regeneration period of 4–5 days was determined to be optimal for the chopped thallus method. The supplementation of amino acids did not have a consistent effect on transformation efficiency, suggesting that ½ Gamborg’s B5 medium alone is sufficient for this method. This finding supports the practicality and simplicity of the chopped-thallus method for large-scale transformations.

Figure 7 illustrates one of the recommended workflows of the chopped-thallus transformation method. While this study primarily used a razor blade, using an electric blender instead can make the process more efficient. In addition, gemmae can be cultured on sterilized cellophane sheets placed over agar plates, allowing the plants to be easily peeled off and collected for chopping. The current transformation efficiency yields approximately six transformants per plant. By using 20 plants as the initial material, it is possible to obtain over 100 transformants. Optimizing the timing of chopping and the conditions for co-culturing could further improve efficiency; however, these challenges remain for future studies.

**Figure 7.**
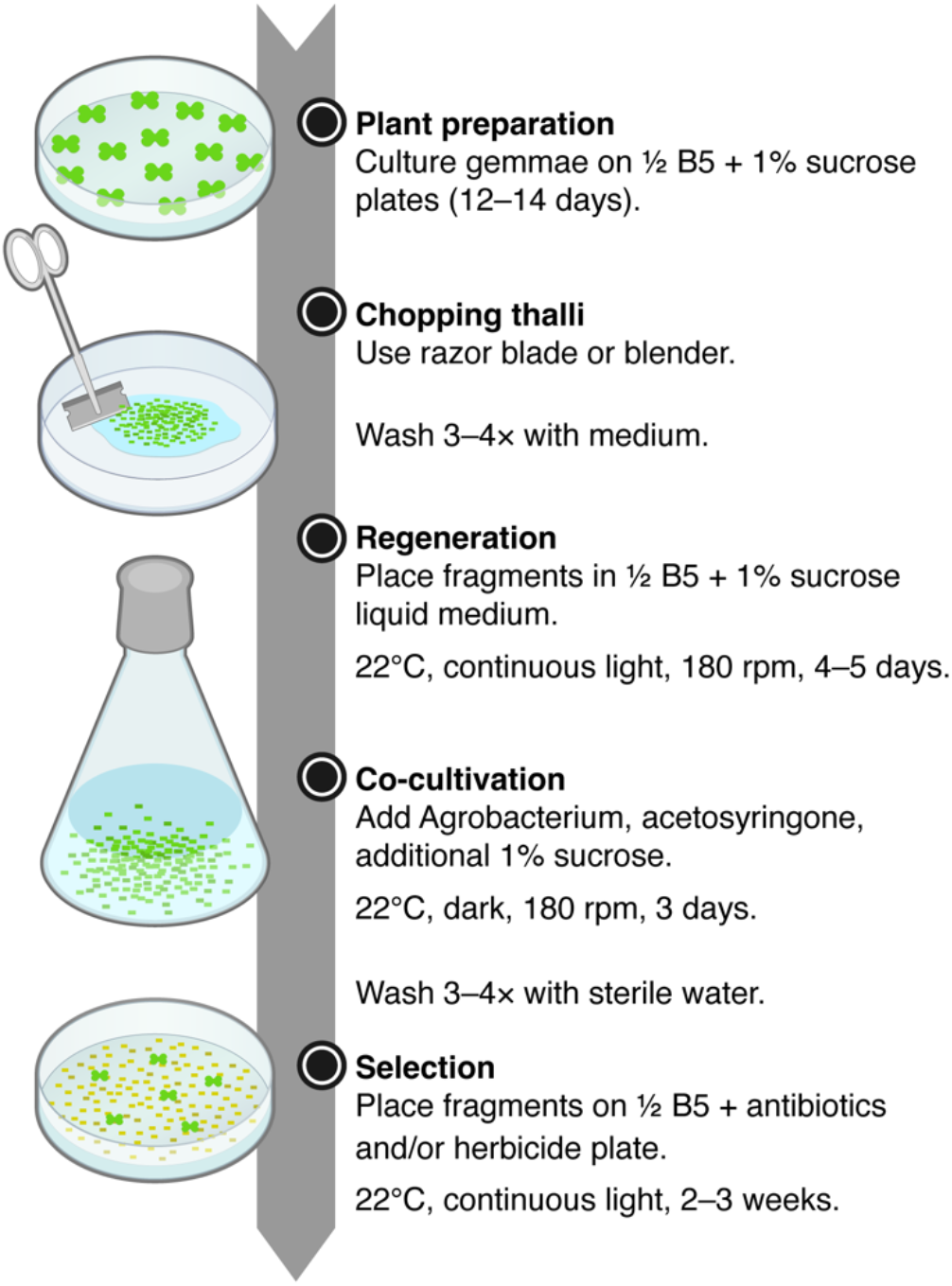
Workflow of the chopped-thallus transformation method.

## 4. Conclusions

The primary objective of this study was to enhance transformation efficiency and simplify the methodology by developing a modified chopped-thallus method. This method uses the entire plant, significantly reducing processing time and maximizing the use of limited plant resources. The chopped-thallus method demonstrated that ½ Gamborg’s B5 medium can effectively substitute for M51C, simplifying the protocol without compromising transformation efficiency. Additionally, supplementation of amino acids to the co-cultivation medium did not provide any significant benefit. The optimal regeneration period for fragmented thalli was estimated to be 4–5 days.

The proposed method is particularly suitable for large-scale transformations, as it eliminates the need to remove meristems or divide each thallus manually, allowing the preparation of numerous thallus fragments simultaneously. This approach facilitates high-throughput experiments, including large-scale mutant screening, by enabling the handling of numerous samples efficiently. Future investigations into the timing of chopping and the conditions for co-cultivation could further optimize transformation efficiency. This simplified and scalable method has the potential to advance genetic studies and support large-scale applications in Marchantia research and biotechnology.

## Author Contributions

Conceptualization: S. G. Y.; investigation: R. B., A. P. and S.G.-Y. ; formal analysis: K. B., E. K., and S. G. Y.; visualization: K. B., E. K., and S. G. Y.; writing—original draft preparation: R. B. and A. P.; writing—review and editing: S. G. Y. All the authors have read and agreed to the published version of the manuscript.

## Funding

This work was supported by the National Science Center, Poland, to S.G.-Y., R.B., and A.P. (UMO-2019/34/E/NZ3/00299).

## Data Availability Statement

Data on the area and regeneration rate of Marchantia thallus fragments are available on Zenodo repository, https://doi.org/10.5281/zenodo.14655757. Other data sets are available upon request from the authors.

## Acknowledgments

The pMpGWB303 plasmid was kindly provided by Dr. Takayuki Kohchi (Kyoto University, Japan) through Addgene (#68631). We would like to thank Dr. Takayuki Kohchi and Dr. Shoji Mano (National Institute for Basic Biology, Japan) for their generous provision of liverwort Tak-1. We also thank Dr. Kenji Yamada (Jagiellonian University, Poland) for valuable feedback.

## Conflicts of Interest

The authors declare no conflicts of interest.

## Abbreviations

The following abbreviations are used in this manuscript:

ANOVA: Analysis of variance
EF1: Elongation factor 1-α
GUS: β-glucuronidase
HSD: Honestly significant difference
LB: Left border
mALS: mutated acetolactate synthase Marchantia *Marchantia polymorpha*
RB: Right border
Tak: Takaragaike
TE: Transformation efficiency

## Notes

https://doi.org/10.5281/zenodo.14655757

### Competing Interest Statement

The authors have declared no competing interest.

